# Microtubule dynamic instability is sensitive to specific biological viscogens *in vitro*

**DOI:** 10.1101/2024.05.27.596091

**Authors:** Arthur T. Molines, Claire H. Edrington, Sofía Cruz Tetlalmatzi, Fred Chang, Gary J. Brouhard

## Abstract

Cytoplasm is a viscous, crowded, and heterogeneous environment, and its local viscosity and degree of macromolecular crowding have significant effects on cellular reaction rates. Increasing viscosity slows down diffusion and protein conformational changes, while increasing macromolecular crowding speeds up reactions. As a model system for cellular reactions, microtubule dynamics are slowed down *in vivo* when cytoplasm concentration is increased by osmotic shifts, indicating a dominant role for viscosity in microtubule reaction pathways. In the cell, viscosity is determined by diverse species of “biological viscogens”, including glycerol, trehalose, intermediate metabolites, proteins, polymers, organelles, and condensates. Here we show *in vitro* that microtubule dynamic instability is sensitive to specific viscogen species, particularly glycerol. We found that increasing viscosity with glycerol or trehalose slowed microtubule growth, slowed microtubule shrinkage, and increased microtubule lifetimes, similar to the “freezing” observed previously *in vivo*. Increasing viscosity with a globular protein, bovine serum albumin, increased microtubule growth rates, as its viscous effects may be balanced against its macromolecular crowding effects. At matched viscosities, glycerol had an outsized effect on microtubule lifetimes, rescues, and nucleation compared to other viscogens. Increasing viscosity did not, however, increase the intensity of EB3-GFP comets, indicating that GTP hydrolysis is unaffected by buffer conditions. We propose that glycerol exerts its distinct effect on microtubule dynamic instability by stabilizing the microtubule lattice after phosphate release. Effects of specific viscogens may modulate many cellular reaction rates within local environments of cytoplasm.

## Introduction

The cytoplasm is a complex environment, both in its composition and its physical properties. In eukaryotes, key components determining the physical properties of the cytoplasm are the concentration of ribosomes (Delarue et al., 2018), the concentrations of carbohydrates, such as trehalose (Persson et al., 2020), and the meshwork created by the cytoskeleton, the membranous organelles attached to it (Heinrich et al., 2021), non-membraneous organelles, and biomolecular condensates (Banani et al., 2017). Central among these physical properties is viscosity, which is a measure of the frictional force between fluid layers in relative motion. Classically, the bulk viscosity of a fluid (*η*) determines the diffusion coefficient of a particle in solution, as *D ∝* 1*/η* (Stokes et al., 1851). Similarly, in the high-friction limit of Kramers’s theory, the bulk viscosity determines the rate constant of a chemical reaction, as *k ∝* 1*/η* (Kramers, 1940). In more recent theory, complex aqueous solutions, including cytoplasm, create solvent microenvironments that are defined by a local microviscosity (Barshtein et al., 1995), and this microviscosity determines the rates of biochemical reactions in the microenvironment (Dill et al., 2011). Solvent friction will slow down the rate of protein conformational changes, e.g., in modified Kramers’s models (Ansari et al., 1992), as different domains move and find local minima in the energy landscape (Bryngelson et al., 1995; Dill et al., 1995; Frauenfelder et al., 1991). Experimentally, the viscosity of cytoplasm affects a wide range of processes, including microtubule physiology (Molines et al., 2022; Shen et al., 2024), response to temperature (Persson et al., 2020), intracellular transport (Nettesheim et al., 2020), protein folding (Sekhar et al., 2014), and RNA folding (Dupuis et al., 2018).

Cytoplasm contains many different species of “biological viscogens” (hereafter: bio-viscogens), which we define as molecules, complexes, and higher-order assemblies that create solvent friction *in vivo*. In *S. pombe*, the viscosity of cytoplasm is highly heterogeneous for nano-particles of a fixed size (Garner et al., 2023), varying over 100-fold within individual cells and roughly 10-fold within a clonal population. Such heterogeneity is not observed in cytoplasmic density, however, which is normally-distributed around an average of 285 mgs/ml (Odermatt et al., 2021). Rather, the source of heterogeneity in viscosity is speculated to be organelle distributions and/or heterogeneity in the local species of bio-viscogens, in terms of their size, charge, hydrophobicity, concentration, and molecular structure. Furthermore, the complex makeup of the cytoplasm impacts molecules differently depending on their size (Luby-Phelps et al., 1986; Shahid et al., 2017). Small molecules like ions, amino acids, and simple sugars diffuse freely while larger structures are confined by the cytoskeletal and organelle meshwork. As the cytoplasm contains species of bio-viscogen that range over 3 orders of magnitude in hydrodynamic radius, their contribution to viscosity and macro-molecular crowding could differentially impact cellular reaction rates.

In order to differentiate the effects of specific bio-viscogens, we used microtubule dynamic instability as a model system for a biochemical reaction that can be directly visualized by microscopy (Walker et al., 1988). With microtubules, protein-protein interactions are converted into measurable changes in polymer length as the αβ-tubulin subunits associate or dissociate from the polymer. Microtubule dynamic instability is also an excellent model system because αβ-tubulin’s conformational cycle occurs on length scales that span two orders of magnitude (0.1 nm — 10 nm). When microtubules shrink, their protofilaments peel outwards, deviating by 10’s of nm from the body of the polymer (Mandelkow et al., 1991; McIntosh et al., 2018). At the intermediate scale, αβ-tubulin dimers oscillate between “curved” and “straight” conformations, which differ by up to 1.4 nm (Brouhard et al., 2014). At the smallest scale, αβ-tubulin can “compact” by 0.3 nm in length (Alushin et al., 2014; R. Zhang et al., 2015), perhaps as part of its GTP hydrolysis cycle. Also at this scale is the formation of lateral bonds between αβ-tubulins, which involves movements of small loops at the lateral interfaces (Chaaban et al., 2018). We hypothesized that different species of bio-viscogen, each with different molecular size and structure, may impact these conformational changes distinctly.

To test our hypothesis, we determined the effects three different bio-viscogens on microtubule dynamics, GTP hydrolysis behavior, and nucleation. We chose the bio-viscogens for their physiological relevance and range of physical size (Fig. 1 A). The smallest viscogen is glycerol, a triol with a hydrodynamic radius of 0.3 nm (Schultz et al., 1961) that is produced by *S. pombe* as a cryoprotectant or in response to stress (Soto et al., 1999). Work in the 1970’s and 1980’s showed that increasing viscosity with glycerol increases microtubule polymer formation in turbidity assays (Keates, 1980), and glycerol has long been used in the purification of αβ-tubulin from brain lysates (Ashford et al., 2006; Shelanski et al., 1967). Slightly larger is trehalose, a disaccharide of two glucose molecules with a hydrodynamic radius of 0.4 nm (Olsson et al., 2020; Schultz et al., 1961) that is also produced as a stress protectant in *S. pombe* (Soto et al., 1999). Such stress responses may be part of a larger process of “viscoadaptation”, in which viscogens are produced to regulate cytoplasmic viscosity (Persson et al., 2020). Last, and largest at 4 nm, is bovine serum albumin (BSA) (X. Zhang et al., 2015), which is close to the size of an “average protein” (Milo et al., 2015) and is often used as a macromolecular crowding agent in reconstitutions *in vitro* (Munishkina et al., 2004; Nettesheim et al., 2020; Shahid et al., 2017).

**Figure 1.**
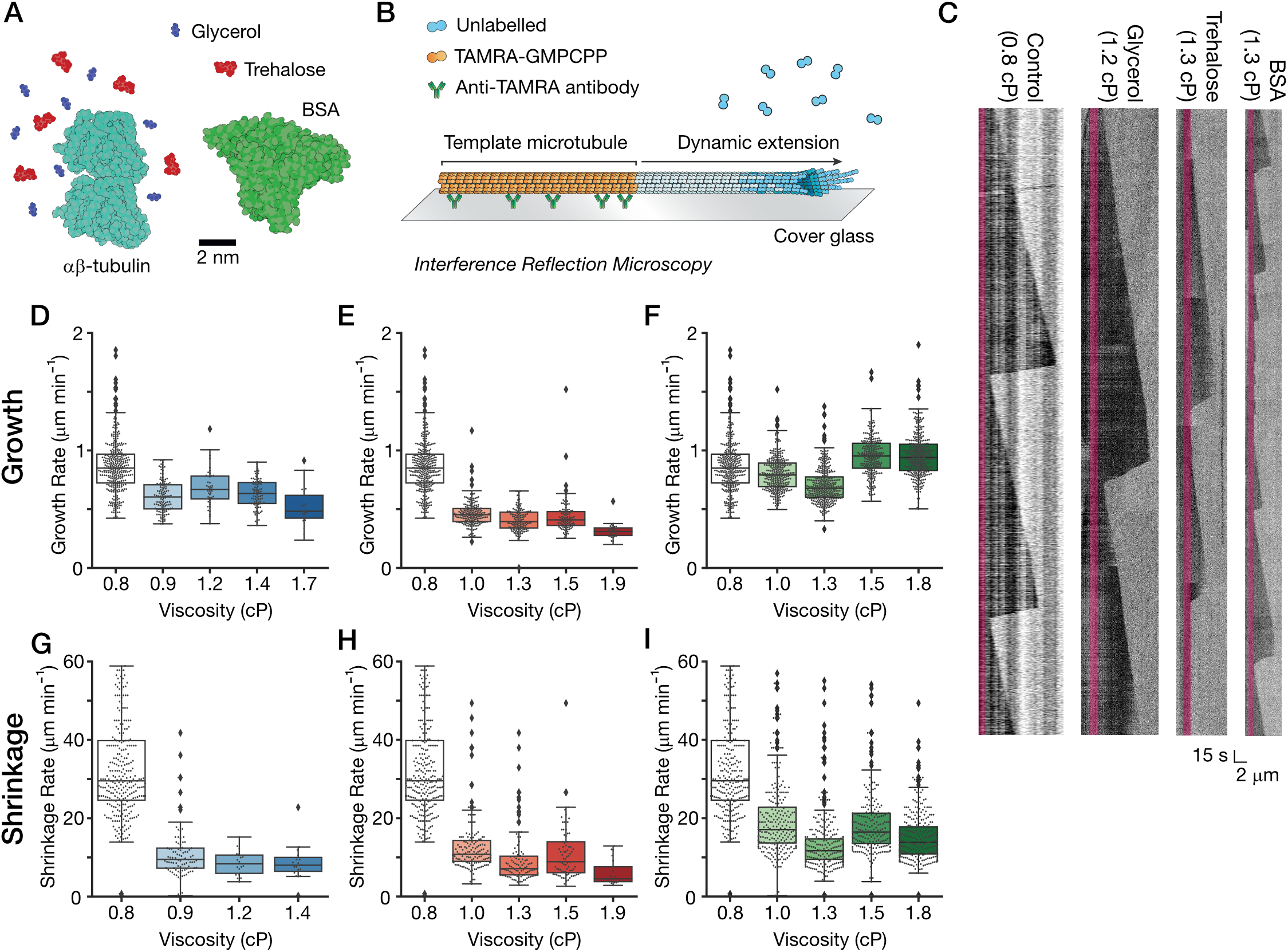
Microtubule growth and shrinkage balance viscous and crowding effects. **(A)** Schematic of the bio-viscogens used in this study drawn to scale. Glyerol (blue), trehalose (red), and BSA (green) were used to increase the viscosity of solutions of αβ-tubulin (teal, labeled). Scale bar 2 nm. **(B)** Schematic of the single molecule assay based on interference reflection microscopy. TAMRA-labeled, GMPCPP-stabilized microtubule tempates are adhered to a cover glass surface using antibodies against TAMRA (see labels). Dynamic microtubule extensions are visualized with IRM. **(C)** Kymographs showing microtubule dynamic instability at 10 *μ*M tubulin in the presence of each viscogen (*η* values indicated). **(D)** Plot of microtubule growth rate versus viscosity for the glycerol titration at 10 *μ*M tubulin. **(E)** Plot of microtubule growth rate versus viscosity for the trehalose titration at 10 *μ*M tubulin. **(F)** Plot of microtubule growth rate versus viscosity for the BSA titration at 10 *μ*M tubulin. **(G)** Plot of microtubule shrinkage rate versus viscosity for the glycerol titration at 10 *μ*M tubulin. **(H)** Plot of microtubule shrinkage rate versus viscosity for the trehalose titration at 10 *μ*M tubulin. **(I)** Plot of microtubule shrinkage rate versus viscosity for the BSA titration at 10 *μ*M tubulin. All data from **(D)** to **(I)** include *n ≥* 3 replicates.

Here we show that the dynamic behaviors of microtubules are sensitive to the specific bio-viscogen species. More precisely, we find that microtubule lifetimes, rescues, and nucleation are significantly altered specifically by glycerol. We propose that glycerol acts by slowing down one or more steps in αβ-tubulin’s conformational cycle. We speculate that complex viscous environments, such as heterogeneous cytoplasm, will impact cellular reaction rates differently depending on the local concentrations of bio-viscogens of varying molecular structure.

## Results

### Microtubule growth and shrinkage are determined by a balance of viscous and crowding effects

To study how dynamic instability is affected by bio-viscogens, we created reaction buffers with matched bulk viscosities ranging from *η* = 0.85 to 1.87 centipoise (cP), as measured using a MEMS-based micro-sample viscometer and temperature-corrected to 32 *°*C (e.g., 20% *m/v* glycerol was measured at *η* = 2.18 *±* 0.01 cP at 23 ^*°*^C and corrected to *η* = 1.67 *±* 0.01 cP, hereafter 1.7 cP). In these buffers, we reconstituted microtubule dynamics using a single molecule assay (Gell et al., 2010) adapted for interference reflection microscopy (IRM, (Mahamdeh et al., 2018)) (Fig. 1B, see Methods). We observed dynamic microtubules that extended from GMPCPP-template microtubules adhered to the cover glass surface as we titrated viscosity. Figure 1C shows an example kymograph for each bio-viscogen (see Table 1 for data). From such kymographs, we measured the full suite of parameters that describe dynamic instability, allowing us to compare these parameters at matched viscosities but with different bio-viscogen species.

**Table 1.** Dynamic Instability Parameters at 10 *μ*M Tubulin.

| [Tb] = 10 μM | CTRL | GLYCEROL | | | | TREHALOSE | | | | BSA | | | |
| --- | --- | --- | --- | --- | --- | --- | --- | --- | --- | --- | --- | --- | --- |
| Viscosity (cP) | 0.85 (±)<br>0.06 | 0.94 (±)<br>0.05 | 1.16 (±)<br>0.02 | 1.39 (±)<br>0.06 | 1.67 (±)<br>0.01 | 1 (±)<br>0.03 | 1.27 (±)<br>0.03 | 1.53 (±)<br>0.01 | 1.87 (±)<br>0.01 | 1 (±)<br>0.07 | 1.32 (±)<br>0.12 | 1.47 (±)<br>0.17 | 1.78 (±)<br>0.17 |
| Number of growth events | 355 | 128 | 45 | 94 | 25 | 207 | 150 | 115 | 34 | 336 | 373 | 279 | 361 |
| Growth rate (μm/min) | 0.86 (±)<br>0.23 | 0.61 (±)<br>0.14 | 0.68 (±)<br>0.16 | 0.63 (±)<br>0.12 | 0.52 (±)<br>0.16 | 0.46 (±)<br>0.11 | 0.41 (±)<br>0.10 | 0.44 (±)<br>0.14 | 0.31 (±)<br>0.06 | 0.80 (±)<br>0.15 | 0.70 (±)<br>0.15 | 0.95 (±)<br>0.17 | 1.04 (±)<br>1.66 |
| Rescue frequency (1/μm) | 0.002 (±)<br>0.002 | 0.028 (±)<br>0.015 | 0.141 (±)<br>0.011 | 0.450 (±)<br>0.137 | N/A | 0.008 (±)<br>0.007 | 0.012 (±)<br>0.011 | 0.016 (±)<br>0.018 | 0.044 (±)<br>0.052 | 0.009 (±)<br>0.007 | 0.013 (±)<br>0.002 | 0.015 (±)<br>0.003 | 0.015 (±)<br>0.001 |
| Number of shrinking events | 301 | 92 | 18 | 18 | 0 | 166 | 114 | 82 | 21 | 301 | 341 | 253 | 331 |
| Shrinkage rate (μm/min) | 35.02 (±)<br>19.11 | 11.67 (±)<br>8.50 | 8.59 (±)<br>3.14 | 11.50 (±)<br>13.99 | N/A | 13.05 (±)<br>7.42 | 10.45 (±)<br>10.87 | 11.23 (±)<br>7.17 | 6.10 (±)<br>3.22 | 20.67 (±)<br>13.24 | 13.51 (±)<br>7.34 | 19.57 (±)<br>11.40 | 15.53 (±)<br>8.09 |
| Lifetime (min.) | 5.3 (±)<br>3.7 | 7.8 (±)<br>6.4 | 13.9 (±)<br>7.3 | 19.2 (±)<br>8.0 | 23.2 (±)<br>5.0 | 7.4 (±)<br>5.0 | 7.7 (±)<br>6.5 | 8.9 (±)<br>6.3 | 11.8 (±)<br>6.8 | 3.7 (±)<br>3.2 | 3.5 (±)<br>2.5 | 3.7 (±)<br>2.8 | 3.4 (±)<br>2.4 |
| Dynamicity (dimers/sec) * | 39.1 | 25.6 | 21.3 | 17.7 | N/A | 20.7 | 17.6 | 17.0 | 12.4 | 37.6 | 33.9 | 44.9 | 45.1 |
| Prob. to nucleate at 15 min. | N/A | N/A | N/A | N/A | N/A | N/A | N/A | N/A | N/A | N/A | N/A | N/A | N/A |

Previously, we showed that microtubule growth rates decrease when viscosity is increased with glycerol (Molines et al., 2022; Wieczorek et al., 2013). Repeating these experiments at 10 *μ*M tubulin, we observed that glycerol titration decreased the mean growth rate from 0.87 *±* 0.23 *μ*m*·*min^*−*1^ at *η* = 0.9 cP to 0.52 *±* 0.16 *μ*m*·*min^*−*1^ at 1.7 cP (Fig. 1D). Trehalose titrations behaved similarly, decreasing growth rates to 0.31 *±* 0.06 *μ*m*·*min^*−*1^ at *η* = 1.9 cP (Fig. 1E). In other words, a roughly 2× increase in *η* caused a roughly 2× reduction in growth rates. Such inverse proportionality is consistent with a simple two-step model of microtubule growth (Wieczorek et al., 2013) in which αβ-tubulin’s on-rate constant to the end is diffusion-limited, since *D ∝ η*^*−1*^. In contrast, when we increased viscosity with BSA, microtubule growth rates increased modestly to 1.0 *±* 1.7 *μ*m*·*min^*−*1^ at *η* = 1.8 cP (Fig. 1F). This increased growth rate is predicted by the macromolecular crowding effects produced by BSA, which act to increase the biochemical activity of tubulin subunits and can compensate for the increased viscosity (Wieczorek et al., 2013).

Predictions are less clear for microtubule shrinkage. Although shrinkage rates are generally concentration-independent (Walker et al., 1988), and thus should be insensitive to crowding effects, the shrinkage process is complex and multi-state (Luchniak et al., 2023). We observed that all three bio-viscogens slowed down shrinkage rates as they were titrated. These results indicate that solvent friction can slow down the outward splaying of protofilaments (Mandelkow et al., 1991) and/or the diffusional escape of tubulin dimers. The magnitude of this effect differed between bio-viscogen species, however. Glycerol slowed down the mean shrinkage rate from 35.0 *±* 19.1 *μ*m*·*min^*−*1^to 11.5 *±* 14.0 *μ*m*·*min^*−*1^at *η* = 1.4 cP (Fig. 1G). Note: we analyzed the glycerol data at 1.4 cP because too few catastrophes were observed at 1.7 cP to accurately measure the shrinkage rate, as discussed further below, so the effect may be stronger at higher *η*. Trehalose was similar across the titration; shrinkage slowed to 6.1 *±* 3.2 *μ*m*·*min^*−*1^ at *η* = 1.9 cP (Fig. 1H). However, relative to the two small viscogens, the effect of BSA at matched viscosities was more modest, as shrinkage rates slowed to 15.5 *±* 8.1 *μ*m*·*min^*−*1^ at *η* = 1.8 (Fig. 1I). We interpret this effect as evidence that BSA’s macromolecular crowding effects may impact shrinkage in some way.

Our reconstitutions demonstrate the sensitivity of microtubule growth and shrinkage to the molecular nature of the bio-viscogen species, particularly with regards to the balance of viscous effects and crowding effects. Similar complex interactions between viscosity and crowding have been observed for actin filaments *in vitro* (Drenckhahn et al., 1986). These interactions confirm that the molecular nature of bio-viscogens is critical.

### The impact of viscosity on lifetimes and rescues depends strongly on the bio-viscogen

The molecular nature of the bio-viscogen also appeared to affect microtubule lifetimes and rescues. Because very few catastrophes were observed with 10 *μ*M tubulin, we reduced the tubulin concentration to 5 *μ*M. We observed very long life-times in our glycerol treatment, whereas the trehalose and BSA treatments appeared perturbed but closer to control (see Table 2). To characterize these differences, we plotted the cumulative frequency distribution of microtubule lifetimes (Gardner et al., 2011; Odde et al., 1995) and calculated mean life-times. The glycerol titration increased mean lifetimes roughly 3.5× from 4.9 *±* 3.9 min to 16.4 *±* 5.7 min at *η* = 1.7 cP (Fig. 2A). The trehalose titration also increased mean lifetimes but only 2× (8.5 *±* 5.6 min at *η* = 1.9 cP, Fig. 2B). These results suggest that solvent friction can slow down some rate-limiting step(s) in the microtubule catastrophe process, but the molecular structure of glycerol makes it more potent. In contrast, the BSA titration actually decreased microtubule life-times slightly (3.3 *±* 2.5 min at *η* = 1.8 cP, Fig. 2C), suggesting that growth in crowded buffers is destabilizing to the microtubule end. Figure 2D shows all mean lifetimes plotted against viscosity, and the different effects of the three bioviscogens are clear.

**Table 2.** Dynamic Instability Parameters at 5 *μ*M Tubulin.

| [Tb] = 5 μM | CTRL | GLYCEROL | | | | TREHALOSE | | | | BSA | | | |
| --- | --- | --- | --- | --- | --- | --- | --- | --- | --- | --- | --- | --- | --- |
| Viscosity (cP) | 0.85 (±)<br>0.06 | 0.94 (±)<br>0.05 | 1.16 (±)<br>0.02 | 1.39 (±)<br>0.06 | 1.67 (±)<br>0.01 | 1 (±) 0.03 | 1.27 (±)<br>0.03 | 1.53 (±)<br>0.01 | 1.87 (±)<br>0.01 | 1 (±) 0.07 | 1.32 (±)<br>0.12 | 1.47 (±)<br>0.17 | 1.78 (±)<br>0.17 |
| Number of growth events | 154 | 92 | 67 | 26 | 46 | 49 | 63 | 35 | 78 | 195 | 221 | 95 | 227 |
| Growth rate (μm/min) | 0.36 (±)<br>0.09 | 0.43 (±)<br>0.23 | 0.32 (±)<br>0.12 | 0.21 (±)<br>0.05 | 0.27 (±)<br>0.05 | 0.23 (±)<br>0.10 | 0.25 (±)<br>0.07 | 0.19 (±)<br>0.08 | 0.30 (±)<br>0.10 | 0.57 (±)<br>0.15 | 0.64 (±)<br>0.21 | 0.57 (±)<br>0.19 | 0.52 (±)<br>0.11 |
| Rescue frequency (1/μm) | 0.005 (±)<br>0.011 | 0.011 (±)<br>0.18 | 0.261 (±)<br>0.240 | 0.111 (±)<br>0.157 | 0.653 (±)<br>0.566 | N/A | 0.014 (±)<br>0.025 | N/A | 0.031 (±)<br>0.028 | 0.008 (±)<br>0.017 | 0.007 (±)<br>0.005 | N/A | 0.00 (±)<br>0.003 |
| Number of shrinking events | 137 | 69 | 37 | 19 | 9 | 43 | 53 | 34 | 65 | 178 | 185 | 84 | 209 |
| Shrinkage rate (μm/min) | 17.35 (±)<br>8.04 | 11.67 (±)<br>9.25 | 4.73 (±)<br>3.33 | 3.90 (±)<br>4.63 | 4.00 (±)<br>3.23 | 13.94 (±)<br>7.33 | 7.26 (±)<br>3.31 | 6.88 (±)<br>3.93 | 8.43 (±)<br>8.16 | 16.28 (±)<br>11.88 | 15.42 (±)<br>6.79 | 18.12 (±)<br>9.07 | 14.11 (±)<br>6.24 |
| Lifetime (min.) | 4.9 (±)<br>3.9 | 7.5 (±)<br>3.9 | 10.9 (±)<br>6.6 | 12.7 (±)<br>7.0 | 16.4 (±)<br>5.6 | 6.0 (±)<br>4.4 | 6.5 (±)<br>4.1 | 6.2 (±)<br>3.5 | 8.5 (±)<br>5.6 | 3.4 (±)<br>2.6 | 4.6 (±)<br>3.1 | 3.1 (±)<br>2.2 | 3.3 (±)<br>2.5 |
| Dynamicity (dimers/sec) * | 17.0 | 17.9 | 11.1 | 7.1 | 7.5 | 11.3 | 11.1 | 8.6 | 13.2 | 26.8 | 30.6 | 27.1 | 25.2 |
| Prob. to nucleate at 15 min. | 0.56 (±)<br>0.18 | 0.84 (±)<br>0.15 | 0.98 (±)<br>0.01 | 0.89 (±)<br>0.07 | 1 (±) 0 | 0.72 (±)<br>0.11 | 0.77 (±)<br>0.13 | 0.77 (±)<br>0.08 | 0.73 (±)<br>0.16 | 0.60 (±)<br>0.23 | 0.69 (±)<br>0.28 | 0.64 (±)<br>0.14 | 0.61 (±)<br>0.32 |

**Figure 2.**
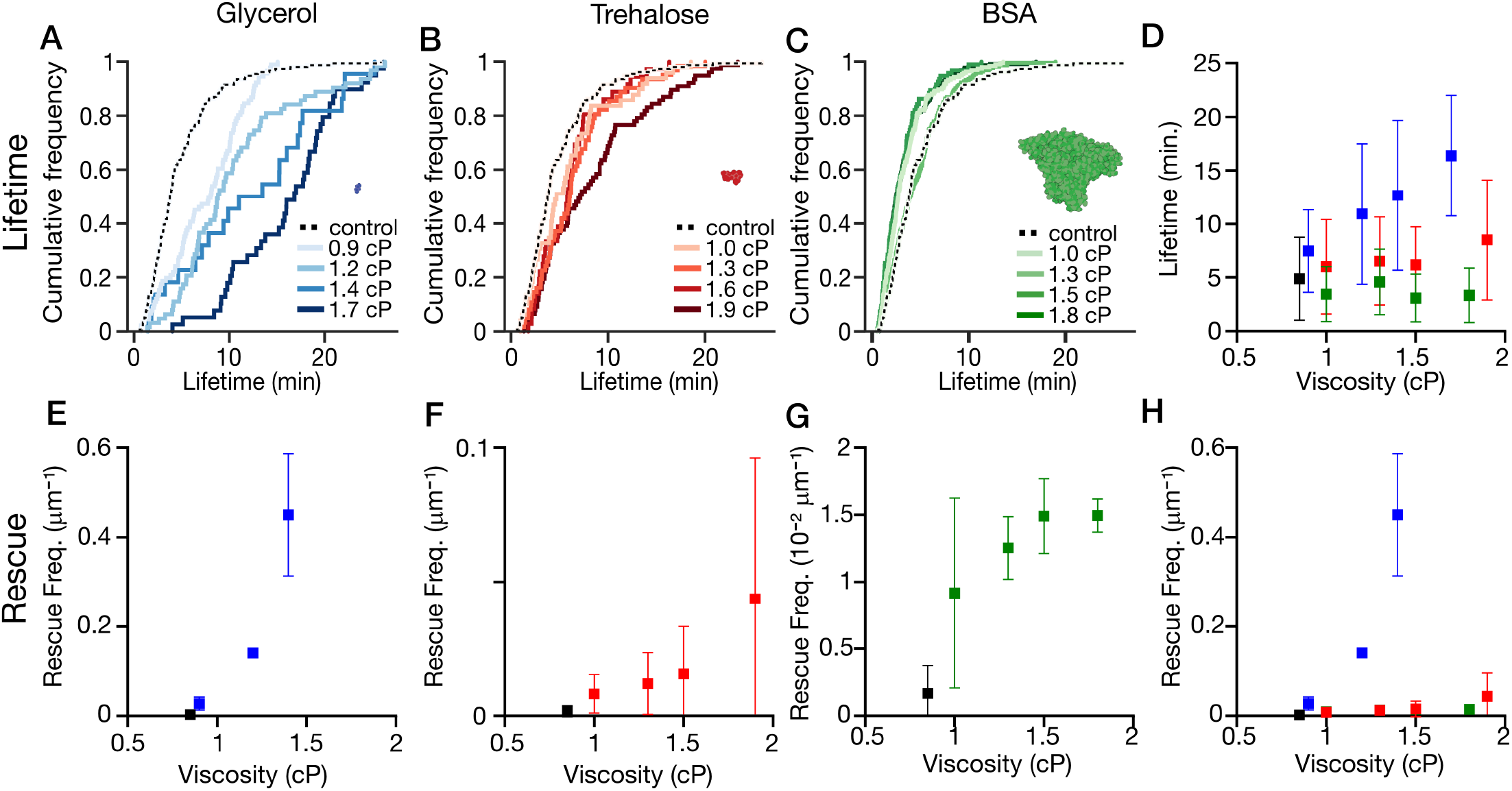
Bio-viscogens impact catastrophes and rescues in distinct ways. **(A)** Plot of cumulative frequency distribution of microtubule lifetimes with glycerol at 5 *μ*M tubulin. Each line represents the total distribution of lifetimes across *n* = 3 replicates. **(B)** Plot of cumulative frequency distribution of microtubule lifetimes with trehalose at 5 *μ*M tubulin. Each line represents the total distribution of lifetimes across *n* = 3 replicates. **(C)** Plot of cumulative frequency distribution of microtubule lifetimes with BSA at 5 *μ*M tubulin. Each line represents the total distribution of lifetimes across *n* = 3 replicates. **(D)** Plot of mean lifetime against viscosity for all three bio-viscogens (blue: glycerol, red: trehalose, green: BSA) at 5 *μ*M tubulin **(E)** Plot of rescue frequency against viscosity with glycerol at 10 *μ*M tubulin. **(F)** Plot of rescue frequency against viscosity with trehalose at 10 *μ*M tubulin. **(G)** Plot of rescue frequency against viscosity with BSA at 10 *μ*M tubulin. **(H)** Plot of rescue frequence against viscosity for all three bio-viscogens (blue: glycerol, red: trehalose, green: BSA) at 10 *μ*M tubulin. All data from **(E)** to **(H)** include *n ≥* 3 replicates.

We observed even greater sensitivity to bio-viscogen structure with microtubule rescues, wherein a shrinking microtubule switches back to growth. We measured rescue frequency as a function of shrinkage length (rescues *· μ*m^*−*1^), following recent work on the subject (Lawrence et al., 2018). All 3 bio-viscogens increased the rescue frequency, but the impact of glycerol on rescue was especially pronounced. At 10 *μ*M tubulin, the glycerol titration increased the rescue frequency by *>*200×, from 0.002 *±* 0.002 rescues *·μ*m^*−*1^ to 0.45 *±* 0.14 rescues *·μ*m^*−*1^ at *η* = 1.4 cP (see Fig. 2E). In comparison, the trehalose and BSA titrations increased the rescue frequency by 20× and 8×, respectively (trehalose, Fig. 2F: 0.04 *±* 0.05 rescues*·μ*m^*−*1^ at *η* = 1.9 cP; BSA, Fig. 2G: 0.015 *±* 0.001 rescues*·μ*m^*−*1^ at *η* = 1.8 cP). Because all of the bio-viscogens cause microtubules to shrink more slowly (Fig.1G–1I), they may provide a larger time window for rescues to occur. But glycerol’s *>*200× effect on rescues must stem from something more significant than giving shrinking microtubules a bit more time. Rather, the effects of glycerol stands out for both rescues and lifetimes, suggesting that its molecular structure allows it to significantly impact the conformational cycle of αβ-tubulin.

### EB comet sizes scale linearly with growth rates regardless of viscous environment

Which steps of the conformational cycle might glycerol impact distinctly? The conformational cycle of αβ-tubulin is coupled to the GTPase activity of β-tubulin. Based on differences between the cryo-EM structures of (mammalian) GTP-lattices and GDP-lattices, obvious candidates are: (1) GTP hydrolysis and phosphate release, which leave behind the unstable GDP-lattice; (2) longitudinal compaction of the αβ-tubulin dimer, in which the dimer shortens by 0.3 nm (R. Zhang et al., 2018); and (3) changes in protofilament twist. These steps may or may not be interdependent, and their sequence remains unclear. But if glycerol slowed any of them down considerably, the microtubule lattice would retain a more stable structure.

We were skeptical, however, that GTP hydrolysis and phosphate release could be affected by solvent conditions, because the GTP in β-tubulin’s exchangeable site is not solvent-exposed in the lattice; rather, the GTP becomes buried when an incoming α-tubulin binds on top of it and completes the hydrolysis pocket (Nogales et al., 1999). To determine if viscosity affects hydrolysis, we estimated GTP cap size using fluorescent EB proteins as a marker for GTP-tubulin (Fig. 3A) (Akhmanova et al., 2008). More specifically, we reconstituted the end-tracking behavior of 30 nM re-combinant EB3-GFP in the presence of our 3 bio-viscogens. At matched viscosities, the EB3-GFP “comets” appeared as very similar, diffraction-limited puncta that grew outward from our template microtubules (Fig. 3B). Using the integrated intensity of these puncta as a readout for comet size, it was immediately apparent that glycerol effects did *not* stand out relative to trehalose or BSA.

**Figure 3.**
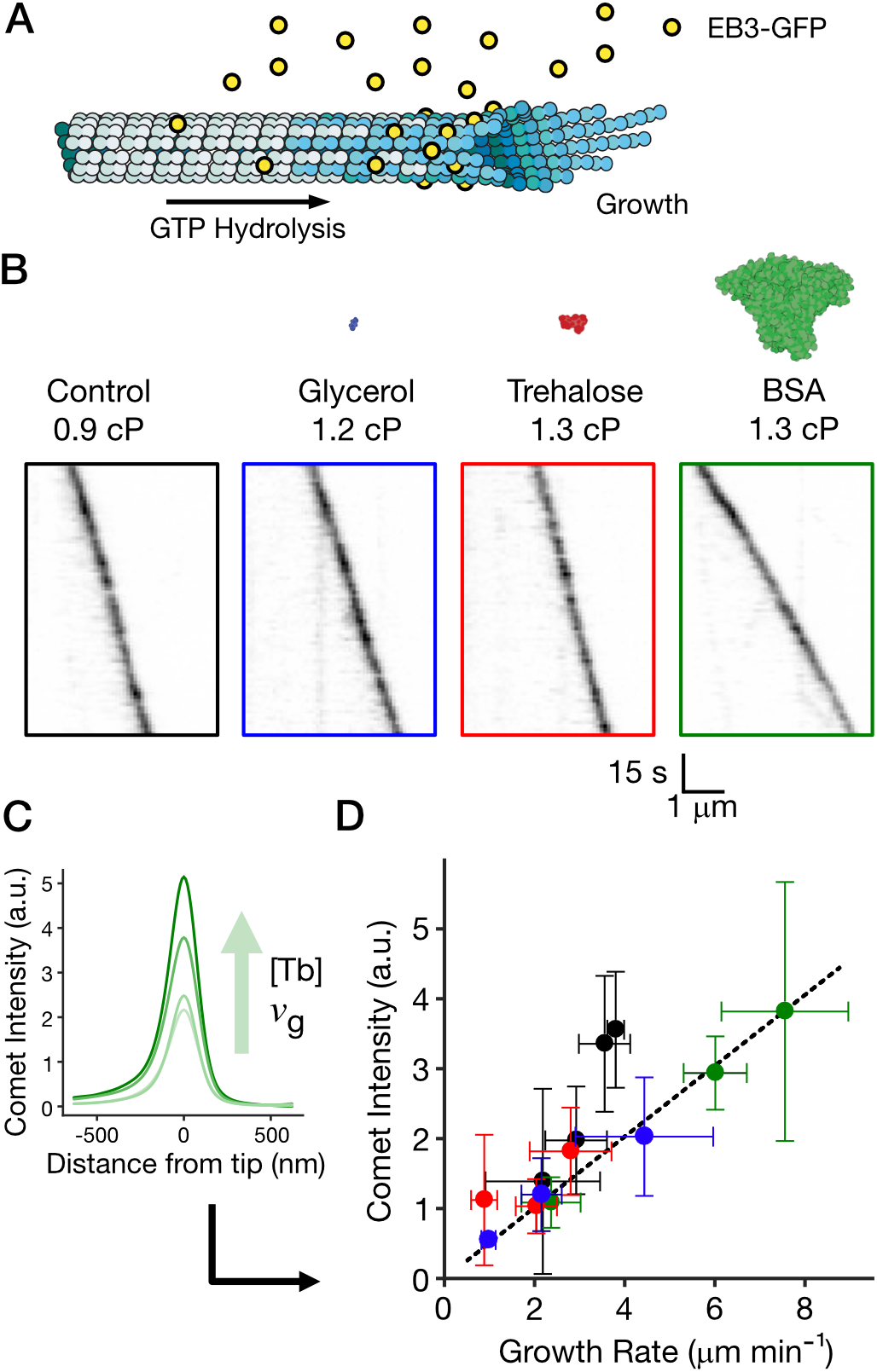
EB comets scale linearly with growth rates regardless of bioviscogen. **(A)** Schematic of a growing microtubule showing the GTP cap (dark blue) and EB3-GFP bindins (yellow) **(B)** Kymographs showing EB3-GFP comets during growth for each bio-viscogen **(C)** Schematic representation of the influence of tubulin concentration and growth rate on comet intensity. **(D)** Plot of comet intensity as a function of growth rate for all three bio-viscogens. Tubulin concentrations are 10 / 20 / 25 / 30 *μ*M for control conditions and 10 / 20 / 30 *μ*M with each bio-viscogen. Data from the three bio-viscogens were fit to a common line (black dashed line).

In reconstituted systems, the mean size of EB comets increases linearly as microtubules grow faster (see inset, Fig. 3C, Bieling et al., 2007), following a simple model that successfully predicts comet shapes (Duellberg et al., 2016). Conversely, EB comets are dimmer when microtubules grow slower. Thus, we predict that comets would become dimmer as microtubule growth became slower as glycerol or trehalose were titrated. Interpretation of such results would be complex, however, because solvent friction may impact EB3-GFP directly, e.g. by reducing its diffusivity. Similarly, crowding effects may increase EB3-GFP’s biochemical activity. Thus, we need an experimental design that allows us to know if the effect of bio-viscogens on comet size is simply a function of growth rate or also a function of buffer conditions.

In order to address the problem, we matched our buffers to a moderate viscosity (*η* = 1.2 cP) and varied the microtubule growth rate independently by changing the tubulin concentration. As predicted, increasing growth rates produced brighter comets in all conditions (Fig. 3D, see plot). This approach enabled us to create a plot of comet intensity versus growth rate for all 3 bio-viscogens at a single matched viscosity (Fig. 3D). Strikingly, the data from all 3 species fell on a common line (Fig. 3D, RGB data). This result suggests that the molecular nature of the bio-viscogen has no influence on the size of the GTP cap, and crowding effects from BSA are not observed. The slope of the common line was lower than a viscogen-free control (Fig. 3D, black), and the comets were slightly dimmer in the presence of a bio-viscogen at each growth rate. We believe the simplest explanation of this reduced brightness is the effects of solvent friction on EB3 itself, e.g. reduced diffusivity. In any case, we can safely conclude that our bio-viscogens do not *increase* the size of the GTP cap, nor do changes in GTP hydrolysis rates explain why glycerol is stronger than trehalose and BSA at stabilizing microtubule lattices. Rather, GTP hydrolysis and phosphate release appear to occur at “normal” rates.

Comparing glycerol to other microtubule stabilizers, we noted that the rescue frequency with glycerol is very close to the rescue frequency in the presence of 400 nM CLASP2*γ* (0.3-0.7 rescues *·* μm^*−*1^, (Lawrence et al., 2018)). Mechanistically, CLASP family proteins cause microtubule lattices to retain “GTP-like” character: CLASP has been shown to maintain microtubule ends in a stabilized intermediate state (Lawrence et al., 2023; Lawrence et al., 2018) and to incorporate fresh GTP-tubulin into sites of lattice damage (Aher et al., 2020). We propose that glycerol’s mechanism of action operates on similar principles. In other words, microtubules stabilized by glycerol remain stable long after phosphate release.

### Spontaneous nucleation and templated nucleation differ in their sensitivity to bio-viscogens

Glycerol’s ability to stabilize lattices after phosphate release may explain its well-known impact on microtubule polymer mass (Keates, 1980) and its usefulness in tubulin purification protocols (Ashford et al., 2006; Castoldi et al., 2003). Alternatively, it’s possible that microtubule polymer mass is sensitive to shrinkage rates, which all bio-viscogens reduce. To distinguish between these possibilities, we performed assays to determine how these bio-viscogens impact microtubule nucleation.

More precisely, microtubule nucleation is divisible into two subtypes: (1) spontaneous nucleation, or the *de novo* formation of stably-growing polymers from soluble αβ-tubulin (Voter et al., 1984), and (2) templated nucleation, or the formation of a stably-growing polymer from a template such as the *γ*-tubulin ring complex, a severed microtubule, or a stable microtubule seed (Roostalu et al., 2017; Wieczorek et al., 2015). Both types of microtubule nucleation are complex processes that must overcome kinetic barriers (Erickson et al., 1981; Kuchnir Fygenson et al., 1995; Roostalu et al., 2017), require a “critical concentration” of tubulin before polymers are observed (Wieczorek et al., 2015), and occur stochastically with time.

To characterize templated nucleation in viscous buffers, we measured the probability that a GMPCPP seed nucleated in the first 15 minutes at 10 *μ*M tubulin (Fig. 4A, following Walker et al., 1988; Wieczorek et al., 2015). When glycerol was titrated, the probability increased from *P* = 0.56 *±* 0.18 to *P* = 1 at *η* = 1.7 cP. The trehalose titration was similar but muted, with *P* = 0.73 *±* 0.16 at *η* = 1.9 cP. In contrast, the BSA titration did not change the probability significantly (*P* = 0.61 *±* 0.32 at *η* = 1.8 cP). The trend between glycerol and trehalose is reminiscient of the trends in microtubule lifetimes: glycerol is the most powerful, trehalose less so. Thus, templated nucleation and microtubule lifetimes may be linked, consistent with previous work showing that templated nucleation is sensitive to catastrophe factors and anti-catastrophe factors (Wieczorek et al., 2015). We speculate that templated nucleation is limited by early catastrophes, which prevent the nascent polymer from forming a persistently-growing microtubule end.

**Figure 4.**
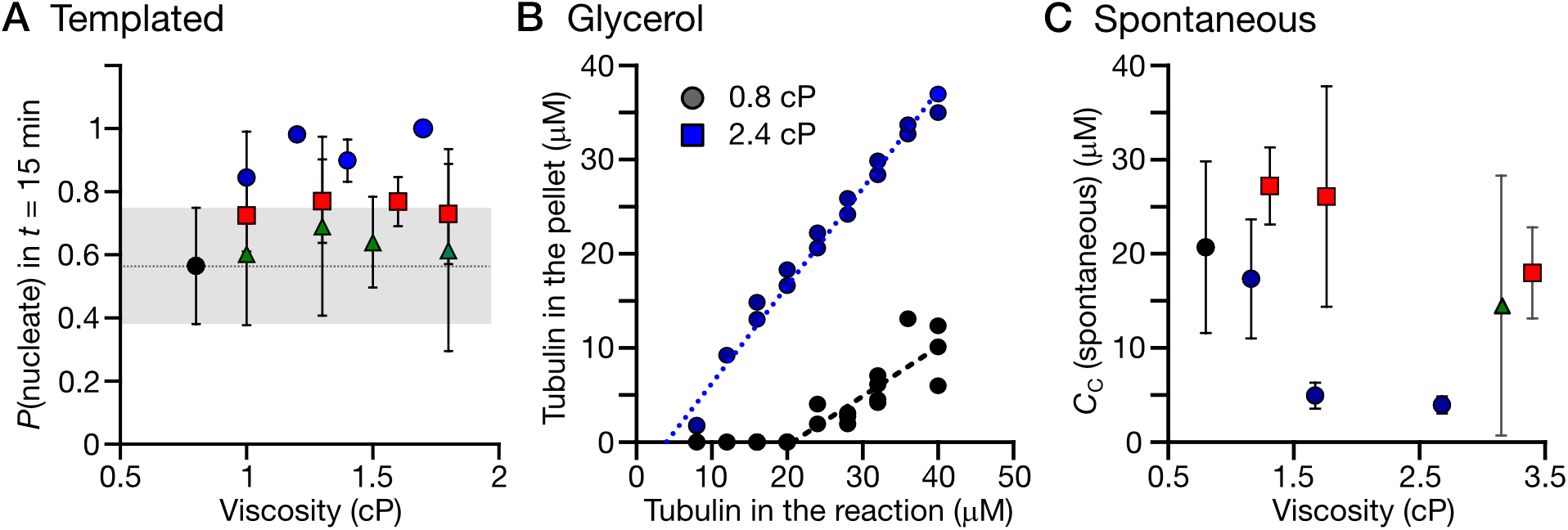
Microtubule nucleation in the presence of bio-viscogens. **(A)** Templated nucleation: Plot of the probability that a microtubule template nucleated a microtubule within a 15 min time window at 5 *μ*M tubulin in the presence of 3 bio-viscogens (glycerol: blue; trehalose: red; BSA: green). **(B)** Spontaneous nucleation with glycerol: plot of the tubulin signal in the pellet versus the total tubulin concentration with glycerol (blue) and control (black) **(C)** Spontaneous nucleation: plot of the critical concentration for spontaneous nucleation as a function of viscosity for all three bio-viscogens (glycerol: blue; trehalose: red; BSA: green).

Next, we used a classic microtubule pelleting assay (Mitchison et al., 1984) to determine the critical concentration required for spontaneous nucleation. Using this method, we measured a critical concentration for spontaneous nucleation of 20.7 *μ*M with a 95% CI of 7.7 to 25.2 *μ*M in control buffer at 35 ^*°*^C (Fig. 4B, black), consistent with recent measurements (McAlear et al., 2022). For these experiments, we titrated glycerol further, so as to reach the 33% (*v/v*) condition frequently used in tubulin purifications (*η* = 2.4 cP). At this bulk viscosity, glycerol lowered the critical concentration significantly to 3.9 *μ*M with a 95% CI of 2.2 to 5.5 *μ*M (Fig. 4B, blue), in agreement with original measurements (Keates, 1980). In contrast, trehalose and BSA had reduced effects on the critical concentration for spontaneous nucleation at similar viscosities (Fig. 4C). Here, the trend across species is reminiscent of microtubule rescues, in that glycerol stands out as distinct. Thus, spontaneous nucleation and microtubule rescues may be linked. From the point of view of solvent friction, the key step in spontaneous nucleation may be to rescue a nascent polymer from a catastrophe. Such rescues are irrelevant for templated nucleation, since the template acts as a “rescue site” itself.

## Discussion

Here we used microtubule dynamic instability as a model system to study how different species of bio-viscogens can influence biochemical reactions. We observed that microtubule growth and shrinkage rates are determined by a balance of viscous effects and crowding effects, but we did not observe strong sensitivity to specific bio-viscogens. In contrast, the transitions of dynamic instability, i.e., catastrophe, rescue, and nucleation, were quite sensitive to specific bio-viscogens *in vitro*, with glycerol having the largest effects relative to trehalose and BSA.

Glycerol’s pronounced impact on microtubule lifetimes and rescues can be understood from both kinetic and thermodynamic models. In the kinetic model, glycerol slows down a critical conformational change; more precisely, glycerol’s microviscosity reduces the rate constant(s) of one or more steps in αβ-tubulin’s conformational cycle (Brouhard et al., 2018) by creating friction along the reaction coordinate or energy landscape. These steps may include the longitudinal compaction of αβ-tubulin dimers (Alushin et al., 2014) or changes in protofilament twist (R. Zhang et al., 2018). If these steps were slowed down, a glycerol-stabilized lattice would retain some structural similarity to “GTP-like” lattices, such as those solved with GMPCPP (Alushin et al., 2014), GTP*γ*S (R. Zhang et al., 2018), GDP-Pi (Manka et al., 2018), and hydrolysis-deficient mutants (LaFrance et al., 2022). We are skeptical that glycerol slows compaction, however, because microtubules trapped in the GTP state bind to EB proteins along their whole length (LaFrance et al., 2022), whereas glycerol-stabilized lattices have dynamic EB comets (Fig. 3). The loss of affinity of EB3-GFP for the glycerol-stabilized lattice indicates that at least some conformational changes, likely compaction, are still occurring.

Other conformational changes beyond compaction and twist are worth considering. For example, some compu-tational models for catastrophes include the formation of “cracks” in the microtubule lattice, or the initial breaking of lateral bonds between protofilaments in regions that may extend beyond the GTP-cap (Li et al., 2014; Margolin et al., 2012; Margolin et al., 2011). Glycerol could reduce such cracks by slowing the movements of loops involved in lateral bonds, namely the M loop and the H2-S3 and H1’-S2 loops. Other computational models explicitly consider the mechanics of αβ-tubulin bending motions, which will be impacted by solvent friction, either in coarse-grained simulations (VanBuren et al., 2005) or molecular dynamics (Fedorov et al., 2019). A Brownian dynamics mechanical model includes an explicit term for viscosity, which enters into Langevin equations that describe the motion of αβ-tubulin within energy potential wells (Zakharov et al., 2015). Regardless of atomic-level mechanism(s), in kinetic models, glycerol acts via reduced rate constants for αβ-tubulin’s conformational changes.

The thermodynamic model considers the entropic component of the energetics of tubulin-tubulin interactions. Like many protein polymers (Oosawa et al., 1975), the energetic drive for microtubule polymerization comes from the release of αβ-tubulin’s shell of structured water and cosolvents; the release increases entropy, or the number of microscopic configurations available to the system (Vulevic et al., 1997). Recently, the thermodynamics of hydrogen bond networks were shown to impact the condensation of proteins with intrinsically-disordered regions, enabling cells to buffer against changes in free versus structured water (Watson et al., 2023). Hydrogen bond networks are disrupted by bio-viscogens like glycerol (Dashnau et al., 2006) and trehalose (Miller et al., 1997), which is why they are used as cryoprotectants. Glycerol’s disruption of hydrogen bond networks may inhibit the reformation of a solvation shell around αβ-tubulin when a microtubule undergoes a catastrophe and collapses. Hence, glycerol may stabilize the microtubule by increasing the entropic penalty to depolymerization.

But why only glycerol? Why can’t trehalose do the same thing at matched viscosities? We can envision three reasons for glycerol’s magic. First, glycerol is the smallest bio-viscogen we tested, with a hydrodynamic radius of 0.3 nm compared to trehalose’s 0.4 nm. Glycerol’s small size has proven useful in studies of protein folding that measure the “effective hydrodynamic radius” (EHR) of protein conformational changes (Sekhar et al., 2013), defined as the average size of the structural elements involved in the conformational change. The EHR can be determined by measuring the dependence of reaction rates on viscogen size (Kalwarczyk et al., 2011; Sekhar et al., 2014) and applying modified Kramers’s equations (Sekhar et al., 2014). Thus, the size difference between glycerol and trehalose may determine their relative ability to affect conformational changes within αβ-tubulin with EHR *<* 0.4 nm. In this first reason, the relevant size scales for the transitions of dynamic instability are quite small and thus sensitive to glycerol (e.g., “cracks” or similar loop movements), while the relevant size scales for polymer growth and shrinkage are large (e.g., diffusion of tubulin dimers and curling of full protofilaments).

Second, glycerol may differentially affect the solvation dynamics and hydrogen bond networks around αβ-tubulin relative to trehalose. Glycerol is a triol and trehalose is a disaccharide, so they interact differently with bulk water. Although both molecules are classified as cryoprotectants, they have different “glass transition temperatures”, an important parameter in cryopreservation studies, and they are frequently combined in glycerol-trehalose mixtures (Cicerone et al., 2004; Curtis et al., 2006). We are skeptical, however, that solvation dynamics can explain glycerol’s distinct effects on microtubule lifetimes and rescues. We note that glycerol and trehalose slow down microtubule shrinkage similarly after a catastrophe (Fig. 2), so the relevant solvation dynamics occur *prior* to αβ-tubulin dissociation and reformation of a full solvation shell. Furthermore, glycerol significantly impacts microtubule lifetimes at only 5% (*m/v*), *η* = 0.9 cP, whereas studies of glycerol’s impact on water structure typically use mixtures on the order of 20% (*m/v*) (Towey et al., 2012). Nevertheless, solvation dynamics and the impact of glycerol of the thermodynamics of tubulin-tubulin interactions cannot be ignored.

Third, we cannot exclude the possibility of glycerol binding to tubulin directly in a way that trehalose does not. This direct binding could impact tubulin allosterically, inducing a more stable conformational state. We currently favor kinetic models based on glycerol’s small size, but it’s likely that both kinetic and thermodynamic changes contribute to stability. Determining the relative contribution of these phenomena may be possible by combining molecular dynamics of tubulin-glycerol and/or tubulin-trehalose solutions with coarse-grained simulations.

The three phenomenona above apply broadly to biochemical reactions in cytoplasm. Thus, the viscosity of cytoplasm will be “felt” differently depending on (1) the size scale of the conformational changes involved, (2) the relative importance of solvation dynamics and hydrogen bond networks, and (3) any other viscogen-specific effects. For example, small bio-viscogens will impact the folding of proteins or RNA which involve sub-nm rearrangements of side chains. Large bio-viscogens may not influence these side chain rearrangements, but they will impact processes occurring at larger scales. Intermediate-scale conformational changes are observed, for example, in the allosteric regulation of metabolic enzymes like phosphofructokinase-1 (Webb et al., 2015), and large conformational changes can be seen in the ubiquitous chaperone Hsp90, which switches between a tightly closed conformation and an open V-shape conformation depending on nucleotide binding (Shiau et al., 2006). Viscogens of all sizes may slow the stepping motion of motor domains (Bormuth et al., 2009; Sozanski et al., 2015) or the movement of large cargos through crowded cytoplasm (Nettesheim et al., 2020; Shen et al., 2024).

In cells, increasing cytoplasm concentration by osmotic shock caused microtubules to “freeze” (Molines et al., 2022), indicating that viscous effects dominate over crowding effects *in vivo*. Extrapolating from our *in vitro* results, we propose that acute hyperosmotic shock acts on microtubule reaction pathways primarily through the increased concentrations of small bio-viscogens that act like glycerol. Interestingly, cells are constantly adjusting the contents and organization of their cytoplasm as an adaptive response (Shamipour et al., 2021). The sensitivity to bio-viscogen species observed here and by others (Nettesheim et al., 2020; Sekhar et al., 2013; Sozanski et al., 2015) potentially gives cells an opportunity to regulate their metabolism by altering the amount of large and small viscogens. For example, yeasts have been shown to produce *both* glycerol and trehalose in response to stress (Klein et al., 2017; Mühlhofer et al., 2019; Petelenz-Kurdziel et al., 2013; Soto et al., 1999), perhaps to ensure a broad impact on many cellular processes (Persson et al., 2020).

Overall, cells can adapt the viscosity and crowding of their cytoplasm by controlling the production of bio-viscogens of different sizes (Persson et al., 2020), changing the organization of metabolic enzymes (Lynch et al., 2020), and modulating their cytoskeleton (Shamipour et al., 2021; Shen et al., 2024). Understanding such viscoadaptation will require comprehensive models for the influence of viscosity on cellular reactions at the sub-nm scale.

## Acknowledgments

Thank you for reading our ode to glycerol, which we dedicate to R. Keates from the University of Guelph and his original study on the effects of glycerol on microtubule polymerization kinetics. We thank M. Sébastien and M. Shred for comments on the manuscript. We thank C. Moraes for providing the loan of the viscometer. A. Molines acknowledges a travel grant from The Company of Biologists (JC-STF2003419). C. Edrington acknolwedges a fellowship from an NSERC PGS-D award and a FRQNT fellowship. F. Chang acknolwedges support from the National Institutes of Health (GM141796). G. Brouhard acknowledges support from Natural Sciences and Engineering Research Council of Canada (#RGPIN-2014-03791), the Canadian Institutes of Health Research (PJT-148702), and McGill University.

## Author contributions

Conceptualization: G.J.B, A.T.M., and C.H.E.; methodology: C.H.E, A.T.M., and G.J.B.; formal analysis: C.H.E. and A.T.M.; Script development: S.C.T.; investigations: A.T.M., and C.H.E.; writing: A.T.M., C.H.E., G.J.B., and F.C.; visualization: A.T.M. and C.H.E.; supervision: G.J.B, A.T.M.; project administration and funding acquisition: A.T.M., F.C, and G.J.B.

## Methods

### Tubulin preparation

Tubulin was purified from juvenile bovine brains via cycles of polymerization and depolymerization, as described previously (Ashford et al., 2006). GMPCPP-stabilized microtubule seeds were prepared by polymerizing a 1:4 molar ratio of tetramethylrhodamine (TAMRA, ThermoFisher Scientific) labeled:unlabeled tubulin (Hyman et al., 1992) in the presence of GMPCPP (Jena Biosciences) in two cycles, as described previously (Gell et al., 2010).

### Microtubule reconstitution assay

Dynamic microtubules were imaged in a reconstitution assay from surface-bound, stabilized MT seeds (Gell et al., 2010). Cover glass was cleaned, as previously described (Helenius et al., 2006), with the exception that, after soaking in acetone and sonicating in 50% methanol then 0.5M KOH, cover glass was treated with plasma for 3 min in a plasma oven (Plasm Etch) instead of treatment with piranha solution. Two silanized cover glasses (22 × 22 mm and 18 × 18 mm) were separated by multiple strips of double-sided tape on custom-machined mounts to create channels for solution exchange. Channels were prepared for experiments by flowing in anti-TAMRA antibodies (ThermoFisher Scientific) and blocking with 1% Pluronic F-127 for 20 min. Channels were rinsed three times with BRB80 before flowing in seeds and placing the chamber on the microscope stage, where the objective was heated to 32 ^*°*^C with a CU-501 Chamlide lens warmer (Live Cell Instrument).

Dynamic microtubules were grown from GMPCPP seeds by filling the channel with either 5 or 10 *μ*M tubulin in reaction buffer: BRB80 (80 mM PIPES-KOH [pH 6.9], 1 mM EGTA, 1 mM MgCl_2_) plus 1 mM GTP, 0.1 mg/mL bovine serum albumin, 10 mM dithiothreitol, 250 nM glucose oxidase, 64 nM catalase, and 40 mM D-glucose. Reaction buffer was prepared on ice before being flowed into the channel with a piece of filter paper. Stock solutions of glycerol, trehalose, or BSA in BRB80 were added to the indicated final concentrations. For consistency, a large aliquot of tubulin was thawed on the day of each experiment, sub-aliquoted, and stored in liquid nitrogen. A separate sub-aliquot was thawed for each individual experiment.

### Microtubule dynamics imaging and analysis

Dynamic label-free microtubules were imaged with interference reflection microscopy (IRM) as described previously (Mahamdeh et al., 2018). Microtubule dynamics were analyzed by manually fitting “multi-segmented lines” to kymographs of growing microtubules using FIJI (Schindelin et al., 2012). Each segment measured either a growth event, a shrinking event, or a pause in between. Sorting of these segments and conversion to lengths, times and rates was done with a custom-built Python script.

### Viscosity measurements

Viscosities of BRB80 + glycerol, trehalose, or BSA solutions were measured using an mVROC Viscometer (Rheosense) at room temperature (23 ^*°*^C as measured by the viscometer). Measurements were made at a flow rate of 1000 *μ*l/min for 5 sec for three replicates of each solution. As the experiments with microtubules are performed at 32 ^*°*^C the viscosities measured at 23 ^*°*^C were converted using viscosity tables for glycerol, the measured equation for trehalose (Galmarini et al., 2011), and the viscosity-temperature-concentration relationship for BSA.

### EB3-GFP purification

The coding sequence for EB3 was inserted into a modified pHAT vector containing an N-terminal 6×His-tag followed by a PreScission site and a carboxy-terminal EGFP followed by a Strep-tag II (Bechstedt et al., 2012). Recombinant EB3 was expressed and purified as described with a few minor modifications (Chaaban et al., 2018). Briefly, EB3 was expressed overnight in BL21(DE3) cells at 18 ^*°*^C, then harvested by centrifugation and cell lysis using an EmulsiFlexC5 (Avestin). Then, EB3 was purified through His60 Ni Superflow resin (Takara Bio Inc.) and eluted from the column by cleavage of the His-tag at the PreScission site. His-tag cleavage was achieved by incubating the resin overnight with PreScission protease at 4 ^*°*^C. The cleaved EB3 was then applied to a StrepTactin Sepharose column from which it was eluted with wash buffer (100 mM Tris-HCl, 150 mM NaCl, 1 mM EDTA, pH 8.0) containing 2.5 mM desthiobiotin and 10% glycerol. Protein purity and concentration were assessed by SDS-PAGE, and by absorbance at 488 nm with a DS-11 FX spectrophotometer (DeNovix, Inc.).

### EB comet imaging

To image EB comets, dynamic microtubules were prepared using the standard reaction mix (80 mM PIPES-KOH pH 6.9, 1 mM EGTA, 1 mM MgCl_2_, 1 mM GTP, 0.1 mg/mL BSA, 10 mM dithiothreitol, 250 nM glucose oxidase, 64 nM catalase, 40 mM D-glucose). EB3-GFP was added to this mix at 30 nM along with 50 mM KCl to promote preferential binding to the GTP cap, and as well as 0.1% methylcellulose to minimize fluctuations of growing microtubule ends. To promote protein stability, EB3-GFP aliquots were thawed and kept on ice rather than freezing sub-aliquots and thawing one out for each experiment, as is the case with tubulin. Time lapse images of EB comets were acquired at 1 second intervals by TIRF microscopy. Total internal reflection fluorescence (TIRF) was achieved by coupling a 488 nm laser to an iLas2 targeted laser illumination system (BioVision, Inc.) equipped with 360^*°*^ TIRF. The objective was heated to 34 ^*°*^C with a CU-501 Chamlide lens warmer (Live Cell Instrument). Images were recorded on an Andor iXon + DV-897 EMCCD camera with a pixel size of 160 nm.

### EB comet analysis

EB comets were selected manually by clicking on the position of the comet near shortly after it appears and shortly before it disappears (or before the end of the time lapse). After this manual selection, a Python script was used to generate average intensity profiles for each condition. For each comet, the manually selected points were used to crop the time lapse around the individual comets. Then, the cropped movies were rotated to align the axis of movement of all comets. Each frame of the cropped movies was interpolated by a factor of 6, to allow for sub-pixel alignment of each frame in the movie at the peak sub-pixel. After background subtraction, these aligned comet timepoints were averaged to each other and combined with the averages of all other comets in that movie to generate average profiles for each condition. The total intensity of each averaged comet was calculated from a line profile 3-pixels wide across the center longitudinal axis of the comet. In these experiments, microtubule growth rate was calculated from the distance in time and space between the manually selected comet positions.

### Microtubule pelleting assay

To determine the effect of viscogens on spontaneous nucleation of microtubules, we used a pelleting assay adapted from (Mitchison et al., 1984; Wieczorek et al., 2015). First, tubulin, at a concentration ranging from 4 *μ*M to 40 *μ*M, was incubated with 1mM GTP in 1× BRB80 on ice for 5 minutes. Then, the tubes were transferred to a heating block and incubated at 35 ^*°*^C for 1 hour. The solution was then transferred to tubes for centrifugation in an airfuge air-driven centrifuge (Beckman) at 30 psi at room temperature for 5 minutes. The supernatant was promptly removed, and the pellets were re-suspended in 1× BRB80 and incubated on ice for at least 10 minutes. Samples were diluted 1:1 with BRB80 and mixed with 2× SDS-PAGE loading buffer. Samples were boiled for 10 minutes and loaded onto a 4-12% SDS-PAGE gel (Gen-Script). The gel was run at 140 V for 1 hour, then rinsed with water and stained with Coomassie for 1 hour. Gels were destained in a 10% acetic acid solution overnight. The intensity of the bands was analyzed with the “Gels” analysis tool in FIJI (Schindelin et al. 2012). To determine the critical concentration for spontaneous nucleation, the x-axis intercept was calculated from a line of fit to all points *>* 1 *μ*M.

